# A precision metric for clinical genome sequencing

**DOI:** 10.1101/051490

**Authors:** Rachel L. Goldfeder, Euan A. Ashley

## Abstract

A requisite precondition for the application of next-generation sequencing to clinical medicine is the ability to confidently call genotype at each coding/splicing position of every gene of interest. Current gold standard technologies, such as Sanger sequencing and microarrays, allow confident identification of the genomic origin of the DNA of interest. A commonly used minimum standard for the adoption of new technology in medicine is non-inferiority. We developed a metric to quantify the extent to which current sequencing technologies reach this clinical grade reporting standard. This metric, the rationale for which we present here, is defined as the absolute number of base pairs per gene not callable with confidence, as specified by the presence of 20 high quality (Q20) bases from uniquely mapped (mapq>0) reads per locus. To illustrate the utility of this metric, we apply it across data from several commercially available clinical sequencing products. We present specific examples of coverage for genes known to be important for clinical medicine. We derive data from a variety of platforms including whole genome sequencing (Illumina Hiseq and X chemistry) and exome capture (including medically optimized capture from Agilent, Baylor Clinical Lab, and Personalis). We observe that compared to whole genomes (with ˜30x average coverage), augmented exomes perform far better for known disease causing genes, but less well for other genes and in untranslated regions. Increasing whole genome coverage improves this discrepancy with an average coverage of ˜45x representing the cross over point where performance equals that of exome capture for disease causing genes. A combination of some genome-wide coverage and augmented exon coverage may offer the most cost effective solution for clinical grade genome sequencing today. In summary, this coverage metric provides transparency regarding the current state of next-generation sequencing for clinical medicine and will inform genotype interpretation, technology improvement, and sequencing platform choices for physicians and laboratories. We provide an application on precision.fda.gov (Coverage of Key Genes app) to calculate this metric.

## 1 Introduction

With the rise of precision medicine, next-generation sequencing is becoming more common in clinical practice. A requisite precondition for the application of next-generation sequencing to clinical medicine is the ability to confidently call
genotype at each coding position in every gene of interest. In addition, inclusion of nucleotides relevant for splicing (the dinucleotides at a minimum) is critical for assessment of disruptions of the open reading frame. Current gold standard technologies, such as Sanger sequencing and microarrays, allow confident identification of the genomic origin of the DNA of interest. The current clinical standard for DNA sequencing, Sanger sequencing, is 99.999% accurate [1]. However, Sanger sequencing does not scale (in cost or time) to a full human genome or even exome. Meanwhile, the cost and turnaround time of next-generation DNA sequencing technologies are now within the lower range of many medical tests [2, 3], enabling physicians to incorporate genomic information into medical decisions, such as disease diagnosis and treatment [4–8].

Whole exome sequencing (WES) and whole genome sequencing (WGS) offer different advantages in the question of clinical sequencing. The major advantage of exome sequencing is cost - capturing the ˜2% of the genome that codes for protein prior to sequencing [9] is cheaper than sequencing the whole genome. In addition, WES achieves higher coverage of targeted regions while still producing less total data, and requires fewer computational resources for analysis and storage. However, WES can suffer from reference bias (an enrichment of reads that match with the reference at heterozygous loci due to probe design), which can result in false negative variant calls. Additionally, some regions of the exome are difficult to capture which can result in low or uneven coverage [10, 11]. Some augmented exome products are specifically designed to capture these difficult regions to ensure high coverage [12].

WGS produces more uniform coverage across the genome, and thus, the data is more suited than WES for structural variant and copy number variant analyses. WGS also enables the interrogation of non-coding regions that are important for regulation, such as promoters and enhancers, and play important roles in disease [13, 14]. However, few medical decisions are currently made from knowledge of variants outside of the coding regions.

Clinical tests must be accurate and precise. The current standard for clinical gene test reporting requires high quality genotypes for 100% of coding bases plus the splice dinucleotides. Illumina next-generation sequencing machines have an average raw error rate of ˜0.5% [15]. To achieve a lower rate of error for the final genotype call, each genomic locus is sequenced multiple times and statistical methods are employed to determine the final genotype. Therefore, adequate sequencing depth is required for accurate and complete genotype calls [16, 17].

Existing metrics to describe depth of coverage typically report genome-wide averages and are therefore insufficient for clinical purposes. A typical statement on a clinical report might be”the genome was covered at a mean fold coverage of 30x” or ”90% of each gene was covered at a mean fold coverage of 10x or more”. These global measurements are inadequate for clinical medicine where a confident call at every coding base pair could mean a difference in a critical medical decision. In exome capture, and to a lesser extent genome sequencing, depth of coverage across the sequenced region is highly variable — as much as 100-fold difference [18]. Additionally, normalizing coverage to a percentage belies the enormous diversity in the size of genes (75 to 2,308,000 coding bps). The field is in need of a standard for the adequacy and quality of data defined as relevant for a given locus to allow confident calling. We present a per-gene metric for evaluating short-read next-generation DNA sequencing datasets to address these issues.

The American College of Medical Genetics and Genomics (ACMG) recommends review and reporting of likely pathogenic or pathogenic variants in specific genes with potentially actionable consequences as part of every clinical exome or genome test [19]. This set of 56 genes represents a particularly important focus for a sequencing standards metric. The Clinical Variation Database (ClinVar) and its NHGRI funded partner the Clinical Genome resource (ClinGen) aspires to define the clinical relevance of genes and variants for use in precision medicine. In this study, we evaluated quality-coverage for five clinical sequencing approaches including WGS and three WES platforms. We also compare to a standard (non-clinical) exome sequencing platform. We present data in 3 groups: i) the 56 ACMG genes, ii) all genes with annotations in ClinVar and, iii) all coding genes in RefSeq.

## 2 Materials and Methods

### 2.1 Sequencing approaches and datasets

We obtained sequencing data from one standard exome sequencing platform (Nimblegen (N=2)) and three clinical exome sequencing platforms: Personalis ACE (N=4), Agilent Clinical Research Exome (N=4), and Baylor Clinical Exome (N=3). We also obtained sequencing data from two whole genome sequencing platforms: HiSeq X (including public data from the Garvan Institute (N=2)^1^ and data from Macrogen (N=1)), and HiSeq 2500 (N=4).

Read lengths varied by approach. Nimblegen datasets contained 50 bp paired end reads. Personalis ACE, Agilent Clinical Research Exome, HiSeq 2500 datasets, and Baylor Clinical Exome datasets contained 100 bp paired end reads. HiSeq X datasets contained 150 bp paired end reads [20].

### 2.2 Gene definitions

The genomic coordinates for the 56 ACMG genes, ClinVar genes, and all coding genes were obtained from the RefSeq annotation. Note that only genes on autosomes were considered for the ClinVar and coding gene sets. We examined two regions of clinical interest (1) coding bases plus the splice dinucleotides and (2) all exonic bases (including the UTRs). Note that the coding bases are a subset of all exonic bases.

### 2.3 Methodology

We aligned all sequence data using BWA MEM [21] version 0.7.10 to hg19.We calculated the depth of coverage at each position within the regions of interest using GATK DepthOfCoverage [22–24], version 3.1.1 with parameter mbq set to 20 or 30 to specify minimum base quality (Q20 or Q30) and - mmq set to 1 to specify a minimum mapping quality of 1. The analytical pipeline is available on precision.fda.gov (Coverage of Key Genes app) and github (https://github.com/rlgoldfeder/coverageOfKeyGenes).

**Table 1.** Average Sequencing Yield

| Sequencing Approach | N | Mean (GB) | Read Length(bp) |
| --- | --- | --- | --- |
| Nimblegen Exome | 2 | 3.87 | 50 |
| Personalis ACE Exome | 4 | 12.78 | 100 |
| Baylor Clinical Exome | 3 | 11.18 | 100 |
| Agilent Clinical Research Exome | 4 | 13.46 | 100 |
| HiSeq 2500 Whole Genome | 4 | 124.07 | 100 |
| HiSeq X Whole Genome | 3 | 114.00 | 150 |

## 3 Results

### 3.1 Determining a clinically relevant coverage threshold

We present a metric for evaluating the coverage of medically relevant genes: the absolute number of positions in each gene with depth of coverage lower than a
given threshold (see below).

To determine a clinically relevant coverage minimum, we calculated the theoretical probability of missing a true heterozygous call at various depths of coverage. At a heterozygous position, there are two alleles (x and y), which are each expected to be present in half of the N reads that align to the position. Therefore, p, the probability of observing allele x in a particular read is 0.5. The probability of observing allele x k times can be modeled as a binomial probability mass function:

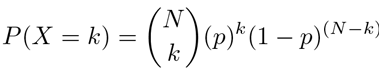

We assume that a heterozygous call would be made if x is present in 20% - 80% of reads. So, the probability of missing a heterozygous call is calculated as:

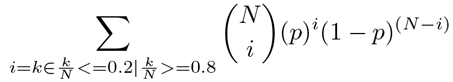

The probability of missing a heterozygous call is displayed in Figure 1 for depths of coverage ranging from 2x to 40x. Based on this distribution, we propose that this threshold should be 20 high quality (Q20) bases from uniquely mapped (mapq>0) reads. We also show results here for a threshold of 30 high quality (Q30) bases from uniquely mapped (mapq>0) reads. Results for other thresholds are available here: https://rlgoldfeder.shinyapps.io/coverage

### 3.2 Total sequencing yield varies by platform

The total yield of sequence data for each sample is described in Table 1. The clinical exomes yielded a mean of 11.18 - 13.46 GB total sequence, the standard Nimblegen exomes yielded a mean of 3.87 GB, and the whole genomes yielded a mean of 111.11 - 124.07 GB.

**Figure 1.**
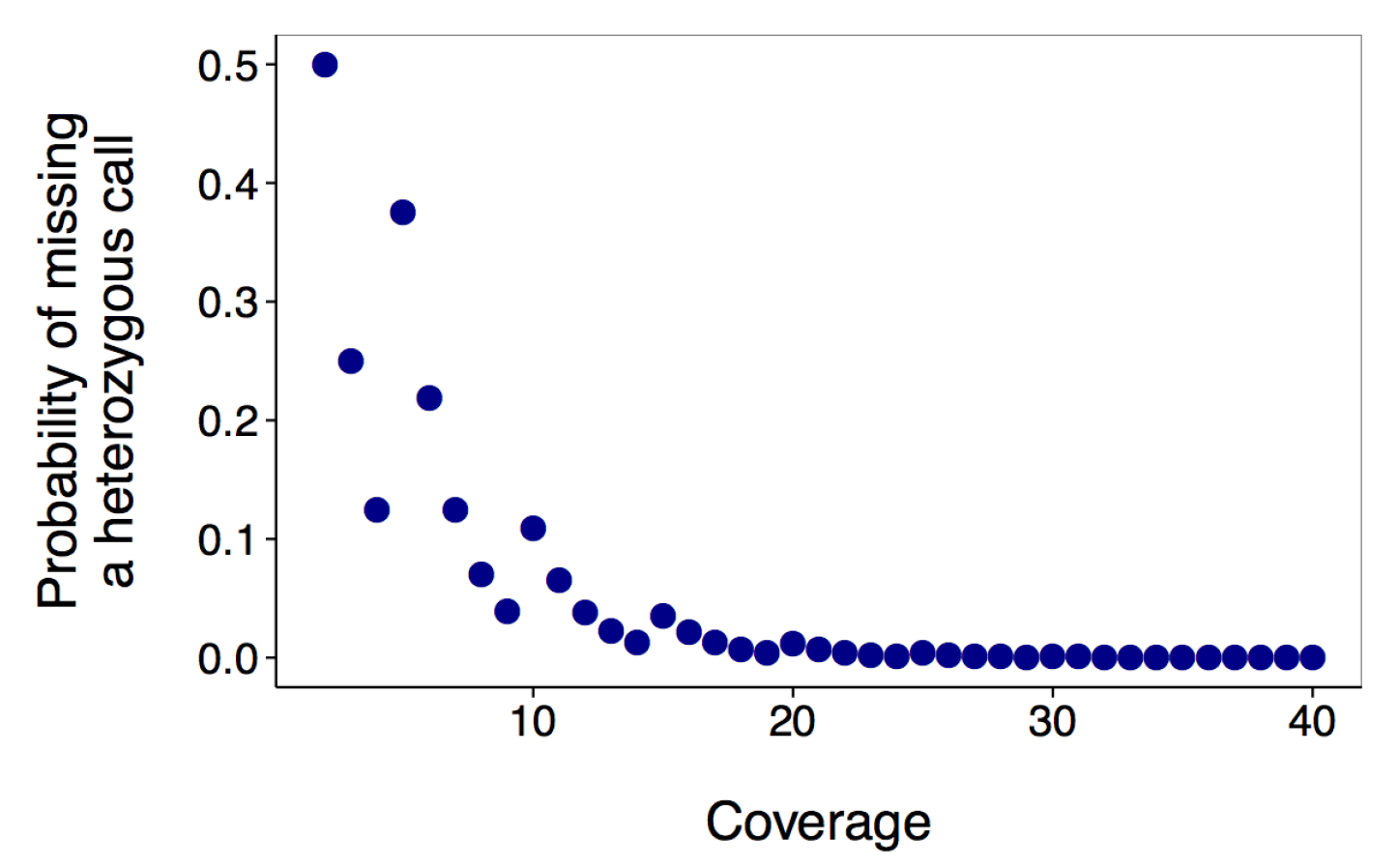
The theoretical probability of miss-calling a heterozygous position at
each depth of coverage ranging from 2x to 40x.

### 3.3 Performance varies across platforms

We calculated the depth of coverage for each coding position in each gene. In the following sections we review results for a single gene, ACMG genes, ClinVar genes, and all coding genes.

#### 3.3.1 Examining KCNH2

Figure 2 shows the coverage of each position in an illustrative (ACMG) gene. KCNH2 is the cause of long QT syndrome type 2, an inherited cardiovascular disease associated with sudden death. We present data with two quality thresholds, Q20 and Q30 (from reads with mapping quality >0) from one sample from each platform. Notably, the whole genome sequencing platforms provide more uniform coverage across this gene, while the exome platforms have more variability. In particular, positions 2,609 - 3,056 (chr 7: 150655144 - 150655591) have low or no coverage for the standard exome and two of the clinical exome sequencing platforms.

**Table 2.** The mean number (and standard deviation) of positions in KCNH2 below the 20 Q20 or a 30 Q30 threshold for each platform.

| Sequencing Approach | Positions Below 20Q20 | Positions Below 30Q30 |
| --- | --- | --- |
| Personalis ACE Exome | 33 (41) | 253 (197) |
| Baylor Clinical Exome | 274 (189) | 413 (187) |
| Agilent Clinical Research Exome | 90 (27) | 122 (49) |
| HiSeq 2500 Whole Genome | 206 (251) | 2,531 (460) |
| HiSeq X Whole Genome | 366 (215) | 2,416 (424) |
| Nimblegen Exome | 1,670 (202) | 2,528 (48) |

**Figure 2.**
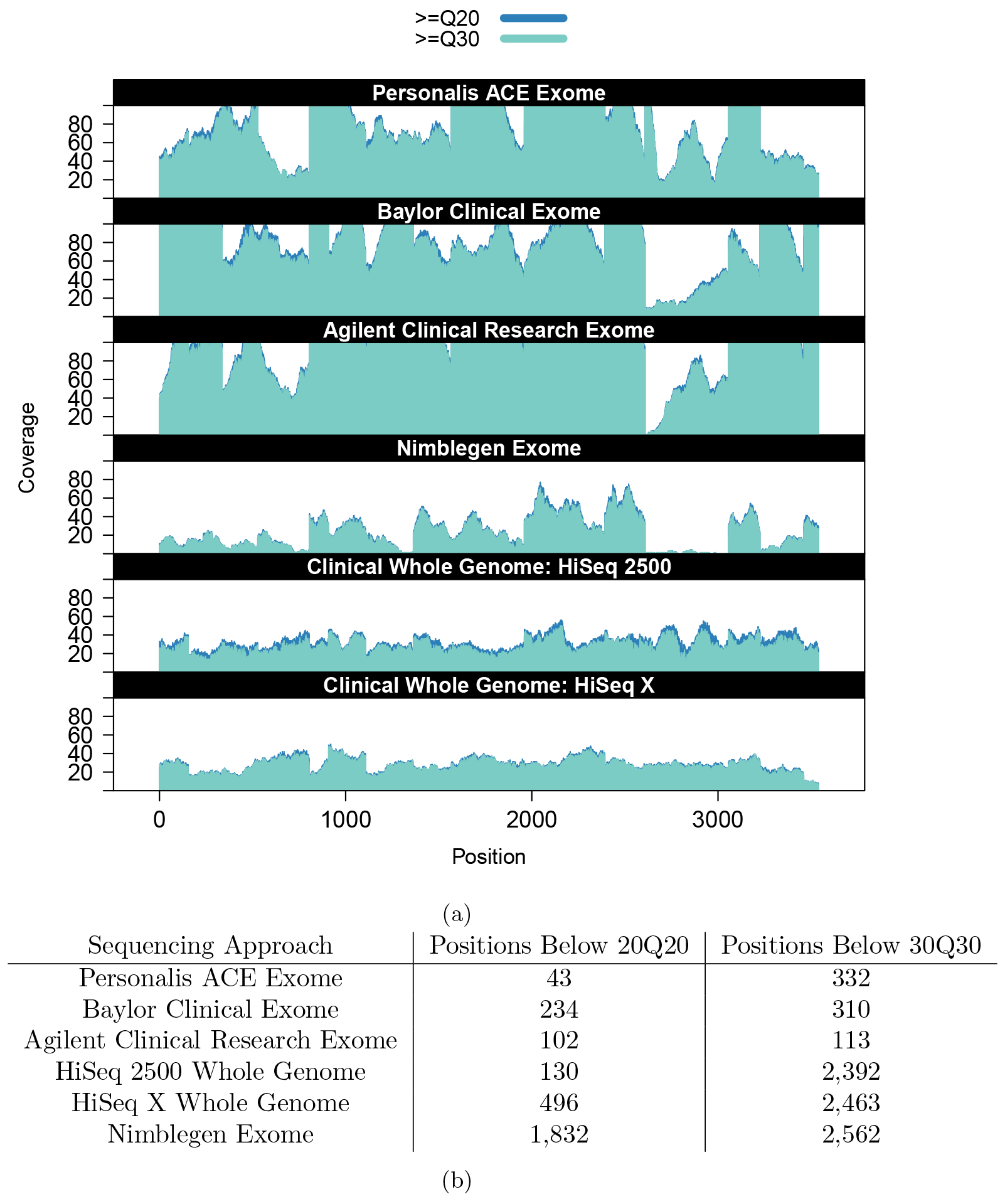
(a) The depth of coverage by minimum Q20 and minimum Q30 bases of each coding position in KCNH2 for one sample per platform (y-axis zoomed in to a max value of 100x). Data plotted in each case derives from the clinically delivered product from each provider. UTRs included in Figure S1. (b) The number of positions in KCNH2 below the 20 Q20 or 30 Q30 coverage threshold for samples displayed in (a).

**Table 3.** The mean number of coding bases in the ACMG gene set failing the
20Q20 or 30Q30 threshold for each platform.

| Sequencing Approach | Positions Below 20Q20 | Positions Below 30Q30 |
| --- | --- | --- |
| Personalis ACE Exome | 805 | 2,457 |
| Baylor Clinical Exome | 2,188 | 3,189 |
| Agilent Clinical Research Exome | 1,288 | 2,242 |
| HiSeq 2500 Whole Genome | 5,377 | 79,316 |
| HiSeq X Whole Genome | 5,800 | 82,311 |
| Nimblegen Exome | 30,353 | 76,281 |

We determined the number of coding loci in KCNH2 that were covered by fewer than 20 Q20 bases or 30 Q30 bases for each sample. Table 2 shows platform means and standard deviations. For the 20 Q20 threshold, the Personalis ACE Exome performed the best (mean positions below 20 Q20 threshold: 33) and the HiSeq X performed the worst of the 5 clinical sequencing platforms (mean positions below 20 Q20 threshold: 366). The Nimblegen standard exome performed the worst (mean positions below 20 Q20 threshold: 1,670). For the 30 Q30 threshold, Agilent Clinical Research Exome performed best (mean positions below 30 Q30 threshold: 122) and the HiSeq 2500 performed the worst (mean positions below 30 Q30 threshold 2,531).

#### 3.3.2 Examining gene sets

##### ACMG genes

We expanded this analysis to all ACMG genes, Figure 3 and Table 3. For these genes, at the 20 Q20 threshold the clinical exome sequencing platforms performed better than the whole genome sequencing approaches, and the standard exome performed worst. The Personalis ACE exome performed best, with a mean of 805 positions below the coverage 20 Q20 threshold for all ACMG genes and the HiSeq X performed worst of the clinical platforms (mean positions below 20 Q20 threshold: 5,800) and Nimblegen standard exome performed worst of all platforms (mean positions below 20 Q20 threshold: 30,353). Several genes consistently have a high (ie: KCNQ1, PRKAG2, RYR1, SDHD) or low (ie: ACTA2, MUTYH, TNNI3) number of bases below the 20 Q20 threshold, while results for other genes (ie: FBN1, PMS2, TGFBR1) were more varied across platforms.

We also examined the number of coding loci in each ACMG gene that failed to meet a coverage threshold of 30 Q30 bases for each sample (Figure S2). In this case, the clinical exome sequencing platforms performed better than the whole genome sequencing approaches. The Agilent Clinical Research Exome performed best (mean: 2,242), and the other clinical exomes performing similarly (means: 2,456 - 3,189). Notably, the number of bases below the 30 Q30 threshold was an order of magnitude higher for the whole genomes and standard exomes (means: 68,117 - 82,311) than the clinical exomes.

Next, we evaluated the number of exonic loci (coding bases plus UTRs) in each ACMG gene below the 20 Q20 and 30 Q30 thresholds, Table 4, Figure S3, and Figure S4. The HiSeq 2500 genomes performed best (mean ACMG gene loci below the 20 Q20 coverage threshold: 9,077), followed by Personalis ACE exomes (mean: 9,299) and HiSeq X genomes (mean: 13,715). The other (clinical and standard) exome platforms performed worse (means: 73,937 - 113,759). Meanwhile, for the 30 Q30 threshold, Personalis ACE performs best (mean: 19,316), followed by the other clinical exomes (means: 77,172 - 82,005), the clinical genomes (means: 110,415 –126,682), and the standard exome (mean: 162,007).

**Fig. 3.**
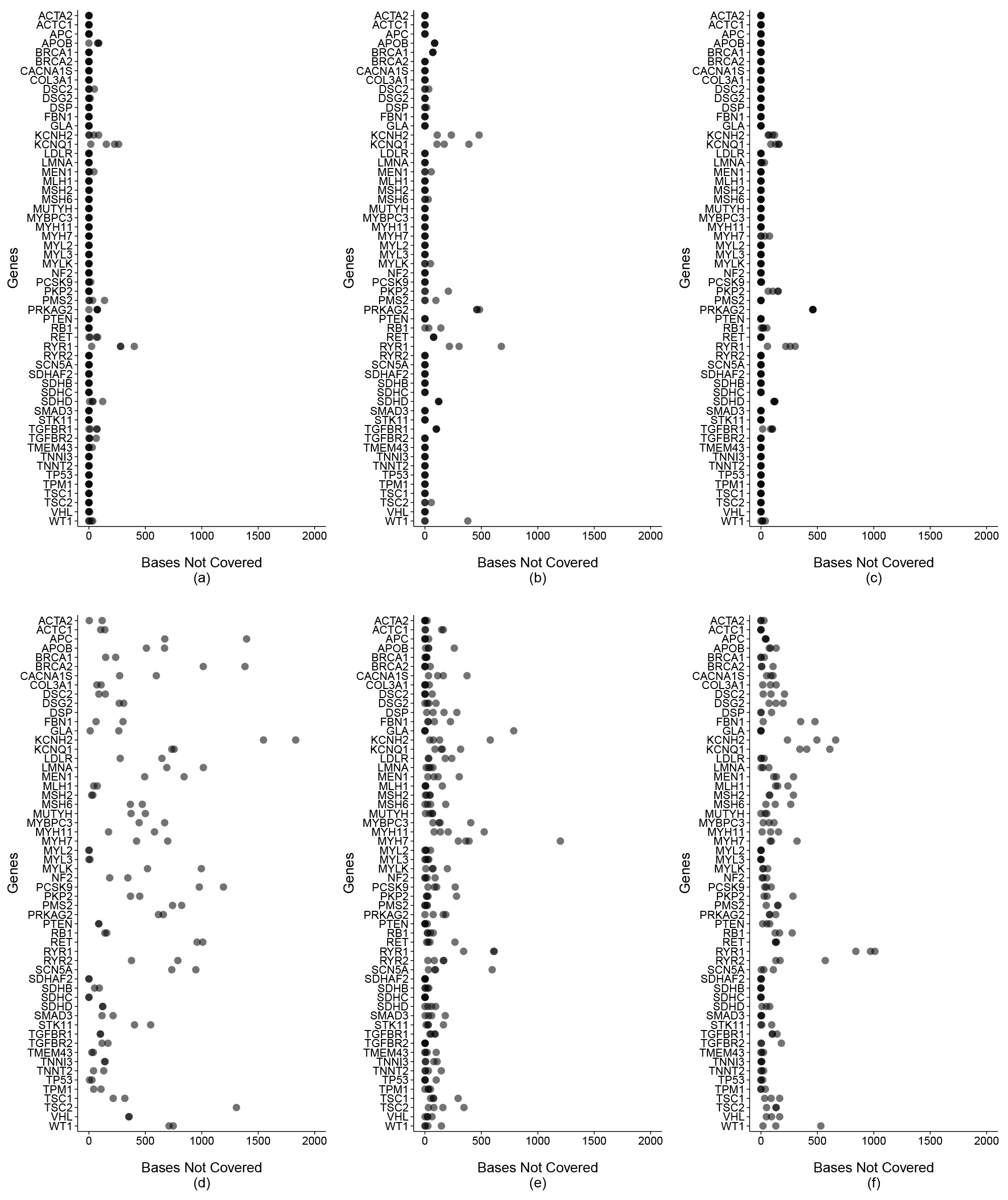
The number of coding loci with <20x coverage by >=Q20 bases from reads with mapQ>0. (a) Personalis ACE Exome (b) Baylor Clinical Exome (c) Agilent Clinical Research Exome (d) Mmblegen Exome (e) HiSeq 2500 Whole Genome (f) HiSeq X Whole Genome. Note that five data points >2,000 omitted from panel (d) and one data point >2,000 omitted from panel (e) for clarity.

**Table 4.** The mean number of exonic bases in the ACMG gene set failing the 20Q20 or 30Q30 threshold for each platform.

| Sequencing Approach | Positions Below 20Q20 | Positions Below 30Q30 |
| --- | --- | --- |
| Personalis ACE Exome | 9,299 | 19,316 |
| Baylor Clinical Exome | 78,235 | 82,005 |
| Agilent Clinical Research Exome | 73,937 | 77,172 |
| Nimblegen Exome | 113,759 | 162,007 |
| HiSeq 2500 Whole Genome | 9,077 | 110,415 |
| HiSeq X Whole Genome | 13,715 | 126,682 |

**Table 5.** The mean number of coding loci below the 20 Q20 threshold for ACMG Genes (N=56), ClinVar Genes (N=3,062), and All Genes (N=18,380) for each platform.

| Sequencing Approach | ACMG Genes | ClinVar Genes | All Genes |
| --- | --- | --- | --- |
| Personalis ACE Exome | 805 | 119,200 | 3,139,544 |
| Baylor Clinical Exome | 2,188 | 159,557 | 2,800,442 |
| Agilent Clinical Research Exome | 1,288 | 167,235 | 2,374,639 |
| HiSeq 2500 Whole Genome | 5,377 | 213,232 | 1,305,055 |
| HiSeq X Whole Genome | 5,800 | 294,598 | 1,803,807 |
| Nimblegen Exome | 30,353 | 1,405,371 | 8,560,901 |

##### ClinVar and RefSeq genes

Next, we examined the coding loci in larger gene sets: i) ClinVar genes and ii) all genes (Table 5 and Figure 5). The augmented exome platforms outperformed the whole genomes approach for ClinVar genes. However, when this analysis was expanded to include genes that currently have no known disease association, the whole genomes performed best. For all three gene sets, the standard exomes performed significantly worse than the genomes or augmented exomes.

### 3.4 Increased overall coverage for WGS improves performance in clinically relevant genes

We showed above that augmented exome sequencing outperforms WGS (with standard sequence yields) for the coding bases of clinically relevant genes. We sought to determine how much sequencing is required for WGS to perform as well as augmented exomes for the coding bases in clinically relevant genes. To do this, we downsampled (using Picard DownsampleSam) one 300x whole genome from the Genome in a Bottle Consortium^2^ to the following average genome-wide coverages: 40x, 45x, 50x, 55x, 60x, 65x, 70x, 75x. Then, for each downsampled dataset, we calculated the number of coding positions that fail the 20 Q20 threshold in each gene set. As overall coverage increases, the number of positions below the threshold decreases. In ACMG genes, WGS performs similarly to the augmented exomes once genome-wide average coverage reaches 45–50x, Figure 6a. The WGS performance in ClinVar genes is similar to the augmented exomes at a genome-wide average coverage between 40x and 45x, Figure 6b. For average coverage WGS levels between 40x and 75x, WGS performs better than augmented exomes, Figure 6c.

**Figure 5.**
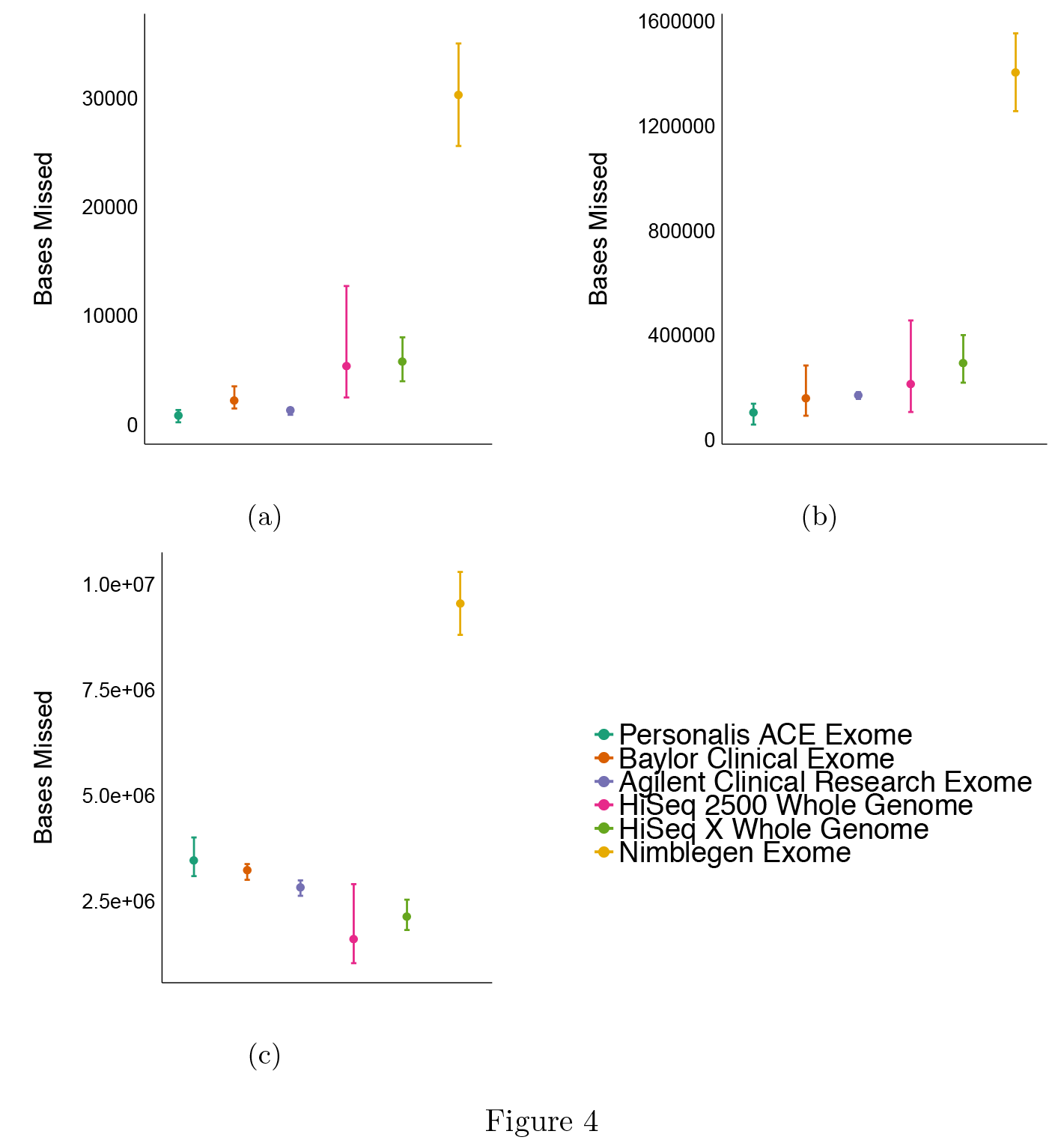
The number of positions below the 20 Q20 threshold (from reads with mapQ >0) for (a) ACMG Genes (N=56), (b) ClinVar Genes (N=3,062), and (c) All Genes (N=18,380) for each sample in each platform. Datapoints represent the mean, bars show the range.

**Figure 6.**
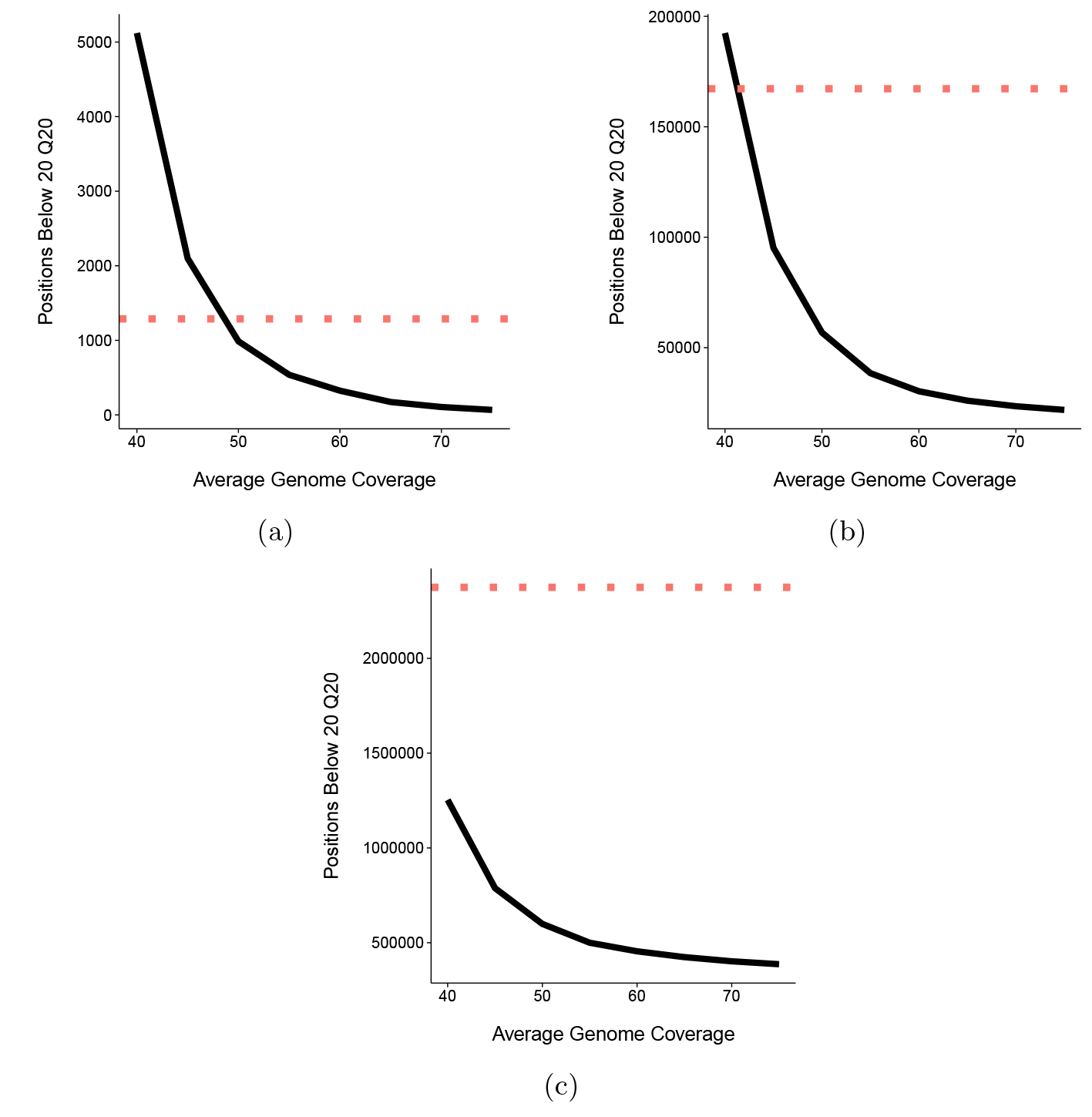
The number of positions below the 20 Q20 threshold (from reads with mapQ >0) for (a) ACMG Genes (N=56), (b) ClinVar Genes (N=3,062), and(c) All Genes (N=18,380) as WGS sequencing coverage increases. Red dotted line shows the mean number of coding positions below the 20 Q20 threshold for the Agilent Clinical Research Exome for comparison.

## 4 Discussion

We present a quality-coverage metric to help evaluate the technical validity of next-generation DNA sequencing for clinical medicine. We use this metric to assess the performance of six sequencing approaches for three gene sets: (1) the 56 most”actionable” genes as described by the ACMG, (2) genes with annotations in ClinVar, and (3) all coding genes.

We observed variability across platforms in the number of loci failing to meet the clinical threshold of 20 Q20 bases from uniquely mapped reads. None of the sequencing approaches met coverage thresholds for all bases in any gene set. The augmented exome sequencing approaches performed better than whole genome approaches in the coding regions of medical genes (ACMG and ClinVar). Meanwhile, the whole genome approaches outperform the augmented exomes across all coding genes. The standard (non-clinical) exome sequencing approach performed much worse in comparison to the clinical approaches.

Notably, when whole genome coverage is globally boosted, performance can approximate the augmented exomes in medically relevant genes. Choice of sequencing approach for a clinical lab will depend not just on effectiveness but also cost-effectiveness. For instance, the right approach may be a combination of low coverage WGS with exon augmentation.

Additionally, we examined the exonic regions (including UTRs) of ACMG genes, and the whole genome approaches outperform two of the three augmented exome approaches. This discrepancy is explained by absence of probes for UTR regions or non-clinical genes in some exome sequencing approaches. For the 30 Q30 coverage threshold, the augmented exomes outperform the whole genomes for the coding and exonic loci in ACMG genes. This is likely due to overall lower coverage of the whole genome sequencing approaches compared to exome approaches.

In clinical practice, variants discovered by next-generation DNA sequencing are still routinely confirmed with Sanger sequencing before results are returned to patients and incorporated into care. False positives are therefore easier to catch and discard, however, it is much more difficult to detect false negatives, which will arise at loci with inadequate coverage. Therefore, identifying areas with inadequate coverage is of the utmost importance. Our metric can be applied to any gene set of interest, and is useful for choosing a platform to use for sequencing or for evaluating the quality of the sequencing results for an individual patient. While minimum coverage thresholds may change as technologies evolve, assessing and reporting the number of bases per gene that fail coverage thresholds will remain vital.

We recently reported that approximately 8% of 100bp reads are not uniquely mapped to the human reference genome [25]. Approaches to handling multiply mapped reads range from excluding the read entirely, to placing the read in all equally-likely locations. Reads that are not uniquely mapped can therefore lead to incorrect variant calls. In our analysis, we included only reads with unique alignments (ie: mapping quality >0), but other applications may require different quality minimums or gene sets different to those presented here. We have made available a browser to interact with the data; users can vary thresholds according to their needs (https://rlgoldfeder.shinyapps.io/coverage). The pipelines to generate this data are also available on precisionFDA (Coverage of Key Genes app) and github (https://github.com/rlgoldfeder/coverageOfKeyGenes) for users to examine their own genes of interest, their own datasets, and any mapping quality, base quality, or coverage thresholds.

## 5 Conclusions

Making a confident call at every coding position of every gene of interest is a sine qua non clinical genetic testing. Confident calling requires adequate read depth and read quality. Here, we present metric to allow comparison across clinical sequencing products with the aim of non-inferiority with current technology. What the clinician needs to know is how many base pairs of the gene of interest are not callable. Thus, the metric describes a sequencing dataset’s coverage as a function of the absolute number of loci with read quality-depth below a given threshold in each gene of interest. As exemplar, we evaluate the coverage of three gene sets by six different sequencing platforms. We hope the methods and results presented will aid clinical sequencing centers in selecting sequencing approaches. Our coverage metric sheds light on the limitations of the next-generation DNA sequencing, which is critical for interpreting results and informing improvements in chemistry and technology.

## 6 Acknowledgments

This work was supported by a National Science Foundation Graduate Research Fellowship and NLM Training Grant T15 LM7033. We thank Amin Zia, Anil Patwardhan, Gill Bejerano, James Priest, and Aaron Wenger for contributing sequence datasets and Daryl Waggott, Jon Bernstein, and Jason Merker for useful discussions.

## Footnotes

1 https://dnanexus-rnd.s3.amazonaws.com/NA12878-xten/mappings/NA12878J_HiSeqX_R1.bam and https://dnanexus-rnd.s3.amazonaws.com/NA12878-xten/mappings/NA12878D_HiSeqX_R1.bam

2 From ftp://ftp-trace.ncbi.nlm.nih.gov/giab/ftp/data/NA12878/NIST\_NA12878\_HG001\_HiSeq\_300x/NHGRI\_Illumina300X\_novoalign\_bams/HG001.hs37d5.300x.bam

## 9 Supplement

**Figure S1.**
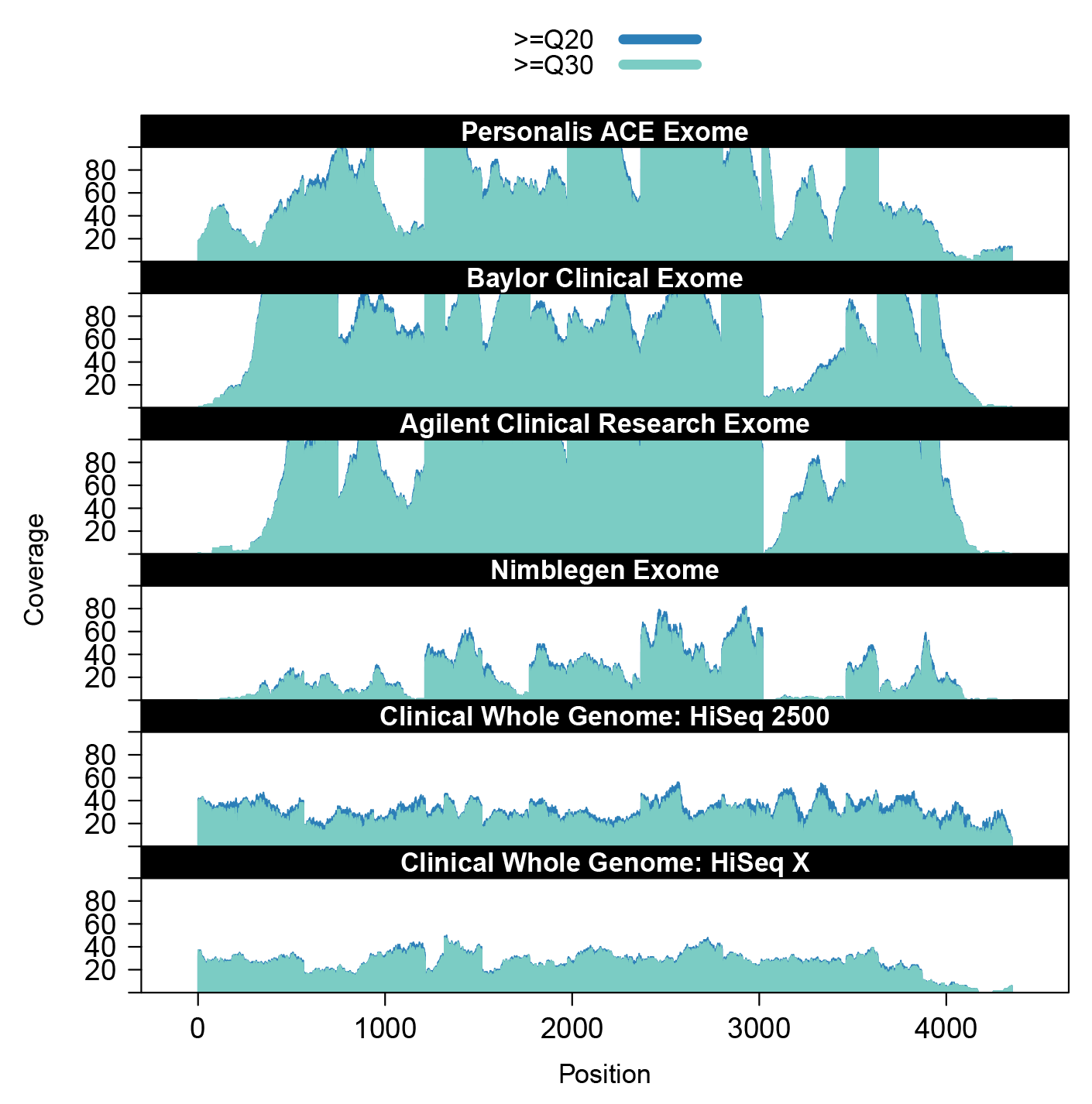
The depth of coverage by minimum Q20 and minimum Q30 bases of each exonic position (including UTRs) in KCNH2 for one representative sample from each platform (y-axis zoomed in to a max value of 100x).

**Figure S2.**
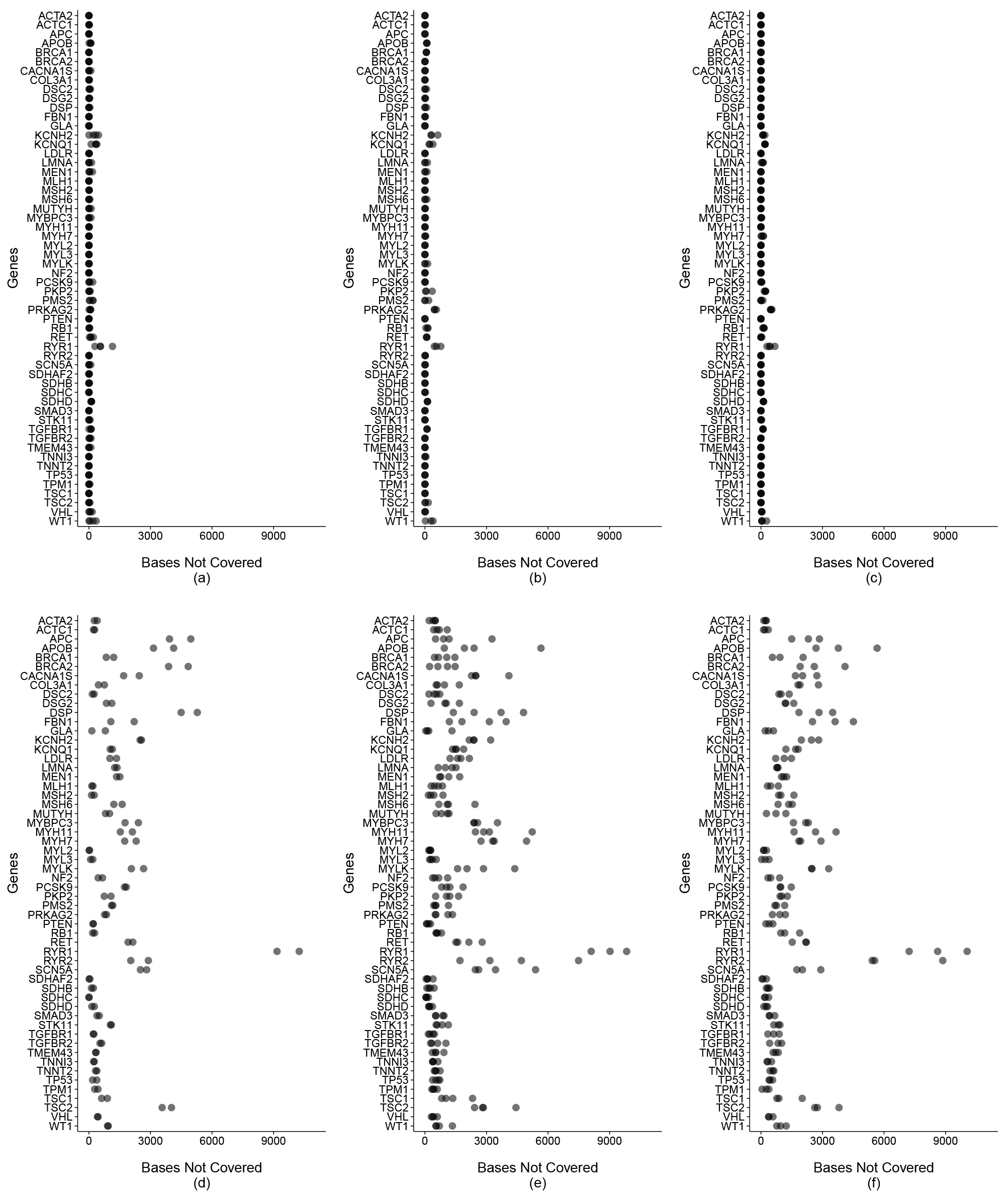
The number of coding loci with <30x coverage by >=Q30 bases from reads with mapQ>0. (a) Personalis ACE Exome (b) Baylor Clinical Exome (c) Agilent Clinical Research Exome (d) N;Lmblegen Exome (e) HiSeq 2500 Whole Genome (f) HiSeq X Whole Genome. Note that one data point >11,000 omitted for clarity from panel (e).

**Figure S3.**
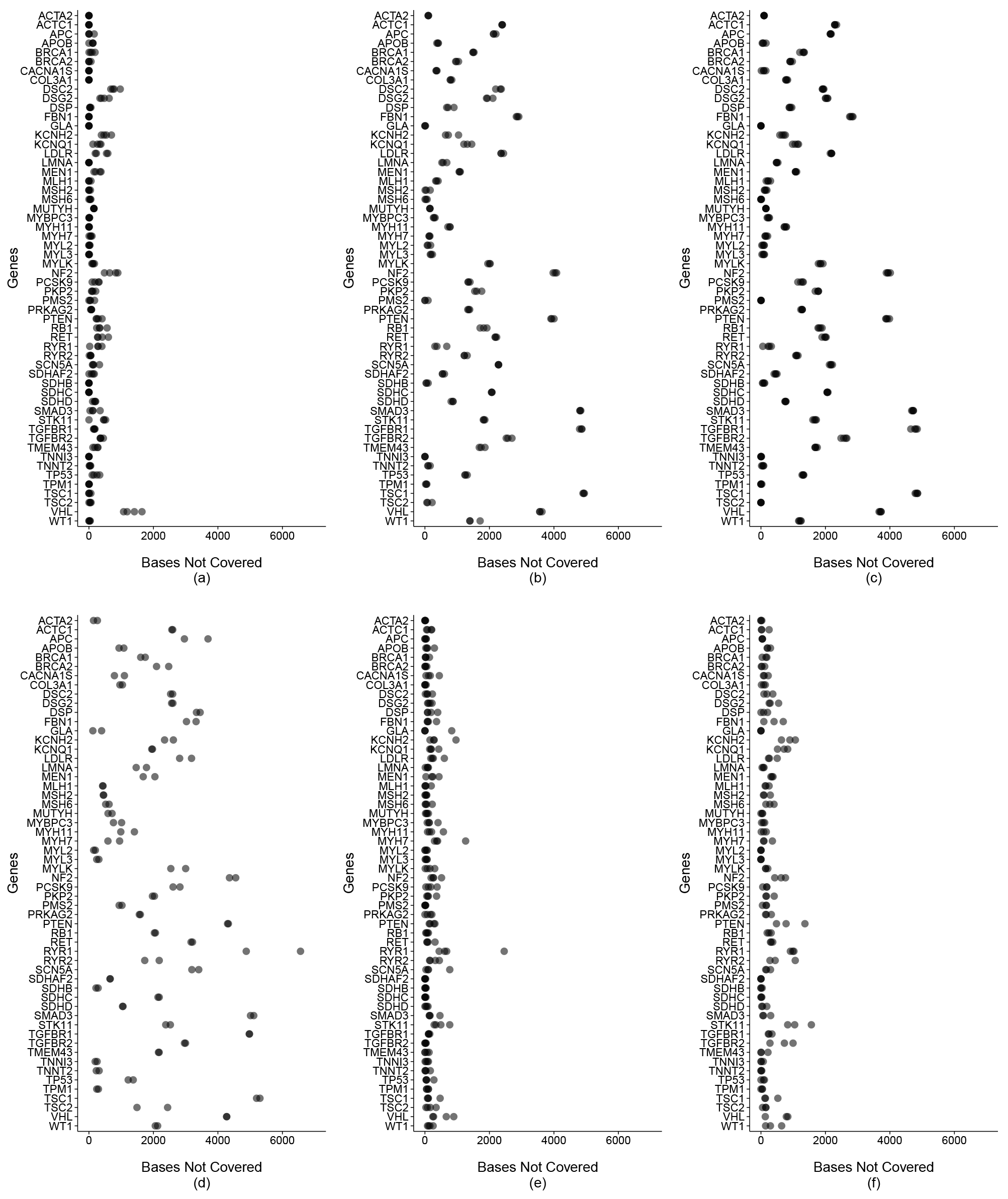
The number of exonic loci with <20x coverage by >=Q20 bases from reads with mapQ>0. (a) Personalis ACE Exome (b) Baylor Clinical Exome (c) Agilent Clinical Research Exome (d) Nimblegen Exome (e) HiSeq 2500 Whole Genome (f) HiSeq X Whole Genome.

**Figure S4.**
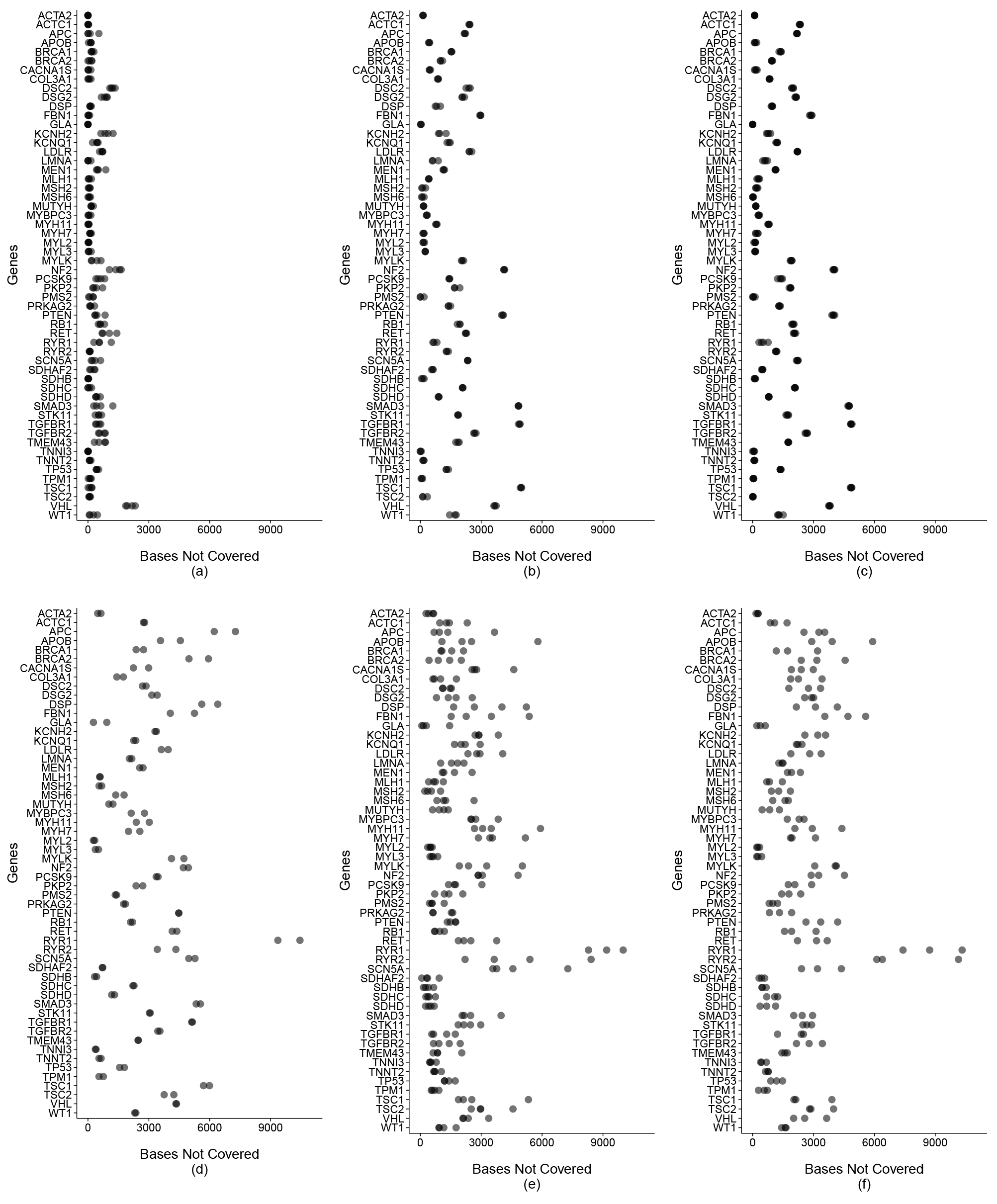
The number of exonic loci with <30x coverage by >=Q30 bases from reads with mapQ>0. (a) Personalis ACE Exome (b) Baylor Clinical Exome (c) Agilent Clinical Research Exome (d) NJimblegen Exome (e) HiSeq 2500 Whole Genome (f) HiSeq X Whole Genome. Note that one data point >11,000 omitted for clarity from panel (e).

